# Iron limitation differentially affects viral replication in key marine microbes

**DOI:** 10.1101/2024.07.22.604475

**Authors:** Charmaine C. M. Yung, Rachel L. Kelly, Kathryn M. Kauffman, Brady Cunningham, Amy Zimmerman, Alexandra Z. Worden, Seth G. John

## Abstract

Viral lysis accounts for much of microbial mortality in the ocean, and iron (Fe) is a critical micronutrient that can limit phytoplankton growth, yet interactions between Fe-nutrition and viral lysis are not well known. Here, we present viral infection dynamics under Fe-limited and Fe-replete conditions for three distinct marine microbes, the photosynthetic picoeukaryote *Ostreococcus lucimarinus*, the cyanobacterium *Synechococcus*, and two strains of the heterotrophic bacterium *Vibrio*. Iron limitation of *Ostreococcus* resulted in slowed growth, and a corresponding decrease in viral burst sizes was observed; this is similar to results from studies of larger eukaryotic phytoplankton (Slagter et al. 2016; Kranzler et al. 2021), where reduced viral replication under Fe-limitation is attributed to the viral reliance on host metabolism and replication machinery. For one strain of *Vibrio*, Fe-limitation similarly impacted viral dynamics, increasing the latent period before infected cells burst to release new virus, and reducing the number of infective viral particles released upon viral lysis. Unexpectedly, for another strain of *Vibrio*, Fe-limitation had no discernible effect on viral replication. Furthermore, dynamics of three *Synechococcus* cyanophages was not affected by Fe-limitation of the host, either in terms of latent period or burst size. The results illuminate the extraordinary ability of some marine viruses, particularly cyanophages, to highjack host metabolism to produce new viral particles, even when host growth is compromised. This has implications for marine ecology and carbon cycling in Fe-limited regions of the global ocean.

## Introduction

Phytoplankton play a crucial role in ocean ecosystems, serving as the base of the marine food chain and sequestering carbon into the deep ocean. Their growth is regulated by both ‘bottom-up’ controls (limiting resources such as dissolved nutrients) and ‘top-down’ controls (mortality processes such as predation and viral infection). While these processes are often examined independently, they are increasingly recognized as interconnected. For example, nutrient limitation in phytoplankton not only impacts their growth rate but also the replication of infecting viruses (1). Viruses rely on host metabolism for replication, so the host’s energy production and availability of specific elements necessary for viral particle production affect replication dynamics. Phosphate-limited hosts exhibit reduced viral replication, characterized by a longer latent phase (the time between initial infection and lysis) and a smaller burst size (the number of virions produced by one infected cell), likely due to the high phosphorus requirement for synthesis of viral genomes (2, 3). Similarly, nitrogen (N) limitation negatively impacts viral replication, possibly due to reduced N availability for synthesizing new viral proteins or general impairment of host metabolism (3).

Iron (Fe) is a necessary co-factor for enzymes involved in all aspects of cell metabolism, from photosynthesis, to energy production, to replication. Iron also plays a major role in regulating global marine primary production, with roughly 30% of the global surface ocean considered Fe-limited (4). Thus, interactions between viruses and iron are of considerable interest, particularly under conditions when host cells become Fe-limited to the extent that their growth rate is reduced. Experiments with the alga *Phaeocystis globosa* and the photosynthetic picoeukaryote *Micromonas commoda*-like Clade C (isolate LAC38) showed a longer latent period and smaller burst size under Fe-limiting conditions (5). Similarly, the diatom *Chaetoceros tenuissimus* appeared less susceptible to viral infection in Fe-limited cultures, and viral infection patterns among natural diatoms suggest less effective viral replication in low-Fe ocean waters (6). Additionally, bacterial siderophore transporters, used to acquire Fe, have been proposed as viral entry sites, linking Fe acquisition with viral lysis (7), a hypothesis which has been explored using correlations between gene abundances and Fe concentrations in Fe-limited ocean regions (8).

Here, we report studies of viral infection among three dissimilar marine microbes. *Ostreococcus lucimarinus*, the smallest known free-living eukaryote, reaches higher cellular abundances and is significantly smaller (9) than the eukaryotic phytoplankton previously studied for iron limitation and viral infection (5, 6). Cyanobacteria such as *Synechococcus* are widespread in the ocean, playing a crucial role in carbon cycling and microbial ecology (2). *Vibrio*, in contrast, are heterotrophic bacteria often found on remineralizing marine organic material and in coastal areas (10). The three host groups selected are not only broadly distributed, but also present in the same eastern North Pacific (ENP) Ocean region (9, 11; Supplemental Data Table 1) where periodic iron limitation has been reported (12, and refs therein). To our knowledge, these are the first studies exploring the interaction between host Fe-limitation and viral infection for these specific phytoplankton host-virus pairs, or in marine bacteria as a whole.

## Results

Iron-replete and Fe-limited cultures of *Ostreococcus lucimarinus* CCMP2972 were separately infected with prasinoviruses OLV1 and OLV7, isolated from the ENP (13). Reduced iron availability resulted in a 43% lower growth rate after acclimation relative to the replete treatment (Table 1). Virus was added to infect both Fe-limited and Fe-replete cultures and the measured multiplicities of infection (MOI) were near 0.9 and 0.4 for OLV1 and OLV7 experiments, respectively. Active viral infection was observed, as indicated by an increase in free viral particles after a latent period, and a corresponding decrease in the concentration of living cells. The dynamics of viral infection were statistically significantly different in the Fe-replete versus Fe-limited treatments. Specifically, infection spread more rapidly in Fe-replete cultures, as evidenced by a shorter latent period, an increased burst size, a larger increase in viral particle abundances by the end of the experiment, and a lower abundance of living cells at the end of the experiment.

**Table 1.**
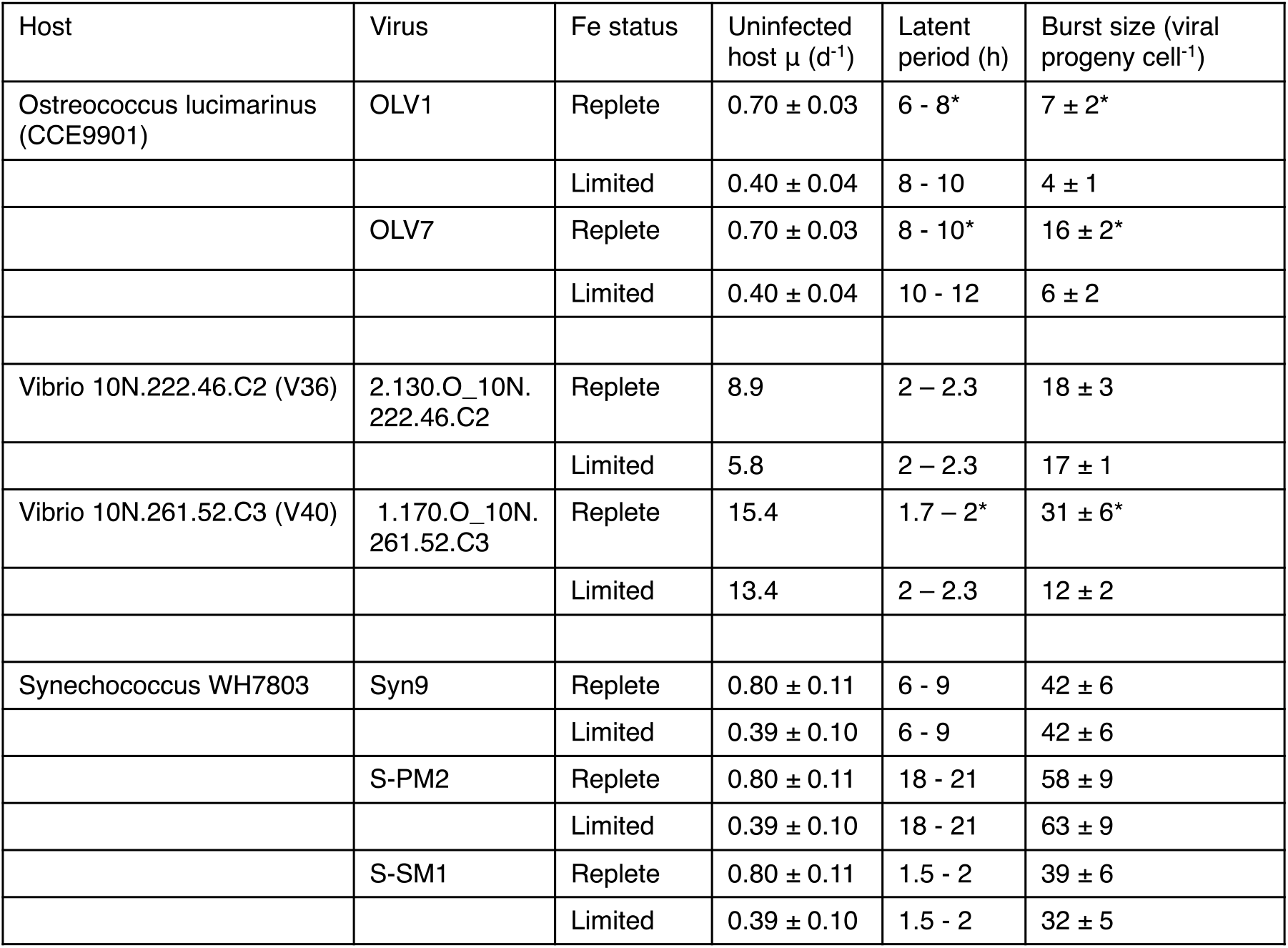
Growth and viral infection dynamics for host-virus pairs. Fe-replete and Fe-limited conditions are determined by diminished growth rates of uninfected host at lower media Fe concentrations, where specific Fe concentrations differ for each host. Latent periods shown as the time period between the last timepoint where a significant increase in free viral particles was not observed, and the first timepoint with a significant increase in free viral particle concentration, and were the same between triplicate experiments. Burst sizes were calculated as the increase in free viral abundance compared to the number of infected cells as described in the Methods, with statistical significance between burst size (*) in Replete versus Limited conditions based on t-tests with *p*-value <0.01.

Parallel experiments were performed with two strains of *Vibrio*, denoted as V36 and V40. The V36 strain was infected with a podovirus, while V40 was infected with a myovirus. These experiments were performed using an MOI of 0.1, and the cells were diluted after the initial infection step, resulting in classic ‘one-step’ experiments tracking a single cycle of viral infection, replication inside the host, and subsequent cell lysis. Iron-limitation reduced growth rates by 35% and 13% for V36 and V40, respectively (Table 1), compared to Fe-replete cultures. The latent period of vibriophage for V36 was not affected by Fe-limitation, and was minimally decreased under low-Fe conditions for V40. There was no change in burst size at the two different Fe concentrations tested for V36 (18 ± 3 in Fe-replete media and 17 ± 1 in low-Fe media), despite growth reduction at the lower Fe concentration. For V40, burst size decreased significantly from 31 ± 6 in Fe-replete media to 12 ± 2 in Fe-limited conditions.

Finally, experiments were performed for *Synechococcus* WH7803 with cyanophages Syn9, S-PM2, and S-SM1, using one-step methodology and an MOI of 0.1. Fe-limited cultures showed growth rate reductions between 45% and 55% relative to Fe-replete cultures (Table 1). Each cyanophage had a distinct latent period and the burst sizes differed among the three cyanophages. When each individual cyanophage encountered the host under low-Fe versus high-Fe growth, the latent period length did not differ significantly. Burst sizes were also similar for each cyanophage under the two host-Fe conditions tested.

## Discussion

The observed reduction in virus replication efficiency under Fe-limited conditions, as observed for *Ostreococcus*, is consistent with studies on larger eukaryotic phytoplankton (5, 6). This result is also consistent with the general tendency for viral replication to slow down when host metabolism is impaired, as for example Fe-deficiencies in humans often negatively impact the ability of viruses to replicate in the host cells (14). Moreover, this result occurred for one *Vibrio* strain as well – indicating marine phages can be restricted in the same manner. However, different from prior reported experiments, our work demonstrates that for some phytoplankton, in this case the cyanobacterium *Synechococcus* and heterotrophic *Vibrio* bacteria (herein strain V36), host Fe-limitation does not impact viral latent period or burst size.

These results point to an extraordinary ability of some marine viruses to highjack host metabolism, in a manner seemingly independent of host-growth rate, akin to recent demonstration of phage replication in dormant hosts (15). The process by which Fe limitation leads to lower cellular growth rates cannot be attributed to a single mechanism, because Fe is involved in so many essential metabolic processes including respiration, photosynthesis, DNA synthesis, and more. While the specific physiological mechanisms underlying Fe-limitation are therefore unknown, diminished growth rate is an unambiguous sign of compromised cell metabolism. It is therefore remarkable that Fe-limitation does not hinder the production of a similar number of virions within the same timeframe, compared to cells that have sufficient Fe and higher growth rates.

Our findings have important ramifications for ocean ecology and carbon cycling, particularly in regions where low concentrations of bioavailable iron limit microbial growth. Viruses capable of effectively replicating within Fe-limited hosts may gain a competitive advantage. This has implications for differentiated host-virus roles in the ‘viral shunt’, a crucial process that redirects both carbon and macronutrients towards recycling within the surface ocean instead of being transferred to higher trophic levels or sinking into the deep ocean (1). The dynamics of viral infection in low-Fe regions may therefore impact how marine microbial communities respond to altered Fe input to the oceans from dust or sediment sources in a changing climate. While more marine phytoplankton and bacterial species and associated viruses should be investigated, the differential interactions between host Fe nutrition and viral infection dynamics demonstrated herein point to possible widespread patterns. If our results reflect generalizable differences in Fe-virus interactions, with viral infection of eukaryotic plankton usually slowing in response to Fe limitation, while viral infection of cyanobacteria remains unaffected by Fe, then they have broad implications for the competition between these two domains of life in low-Fe ocean waters.

## Materials and Methods

For detailed materials and methods please see the SI Appendix.

## Acknowledgments

We are grateful to Camille Poirier for lab support. Funding was provided by the Gordon and Betty Moore Foundation (GBMF3305 to SGJ and AZW), the Simons Foundation (426570SP and LI-SIAME-00001532 to SGJ, and BIO-SCOPE to AWZ) and the Monterey Bay Aquarium Research Institute.

## Materials and Methods

Similar methods were used for each host and virus pair whenever possible, though unavoidable differences in how cells can be cultivated, and how viruses can be propagated and enumerated, led to variations in methodologies as detailed below. Iron concentrations in the media differed greatly between experiments, but in each case were chosen to span a narrow range between Fe-replete conditions, a concentration above which adding additional Fe did not lead to significant increases in growth rate, and Fe-limited conditions, characterized by a significantly lower growth rate compared to Fe-replete conditions. All experiments were performed using trace metal clean techniques; they were grown in polycarbonate or polypropylene labware that had been acid cleaned with 5% HCl and then rinsed at least five times with ultrapure water. All inoculation steps and sampling was performed in class-100 HEPA filtered flow benches. Data are archived at:

### Experiments with *Ostreococcus* and prasinoviruses

#### Initial culturing and viral stock preparation

*Ostreococcus lucimarinus* CCMP2972 (CCE9901) cells (1) were grown in Aquil-based media prepared with artificial seawater, macronutrients, and trace-metal ion concentrations buffered by EDTA (2). The Fe-replete and Fe-deplete media contained 1 μM Fe and 0.01 μM Fe, respectively. The cultures were incubated for 14 days (>10 generations) at 18°C under a 14:10 h light:dark cycle, with a fluorescent light irradiance of 100 μmol photons m^−2^ s^−1^ using semi-continuous batch culturing. Cell density was maintained at 5 ± 0.5 × 10^6^ cells mL^−1^, and growth was monitored using an Accuri C6 flow cytometer (BD Biosciences). Axenicity was confirmed by epifluorescence microscopy after DAPI staining. Additionally, culture samples were tested for bacterial growth after incubation at room temperature in the dark for 1 week following inoculation into Luria broth. Stocks of the prasinoviruses OlV1 and OlV7 (3) with genome features as described in Zimmerman et al. (4), were prepared ‘fresh’ for each experiment from a master stock by infecting exponentially growing CCMP2972 with 1% (v/v) of the master stock and allowing the infected cultures to lyse until clear (∼5 days), with subsequent removal of remaining host cells through gentle vacuum filtration using a sterile Nalgene Rapid-Flow 0.45 μm PES membrane filter. To estimate the proportion of infectious viruses in the concentrated viral lysates, we employed an MPN assay detailed in (4). Briefly, we added 50 μl serially diluted (10^−3^ to 10^−10^) fresh viral concentrate to 150 μl exponentially growing host cells (average daily growth rate over 4 days= 0.75 ± 0.03 day^−1^, target host cell density = 4 × 10^6^ cells mL^−1^) in triplicate 96-well microtiter plates (with 24 replicate wells for each dilution) and incubated them at the above conditions for 14 days. Cell lysis was assessed intermittently for 2 weeks, both visually and by measuring optical density on a plate reader (Molecular Devices SpectraMax 340PC) at 750 nm absorbance. The most probable number (MPN) of infectious viruses was determined based on the proportion of lysed wells using the MPN_ver4.xls Excel spreadsheet (5). Infectivity was calculated by comparing the MPN-estimated abundance of infectious viruses to the abundance of viral particles determined by flow cytometry.

Viral particles were concentrated using VivaSpin20 (Sartorius) 100,000 MWCO PES centrifugal filtration units, then washing each VivaSpin20 unit twice with 0.02-μm-filtered (Whatman Anotop Plus) 1X TE buffer (pH 8.0) and gently vortexing to dislodge additional virus particles. Triplicate 1 ml samples of each concentrated viral lysate was fixed with glutaraldehyde (EM grade, 30 min at 4°C, 0.25% final concentration) and flash frozen. The remaining concentrated viral lysate was stored in the dark at 4°C until used in infection experiments.

#### Quantification of concentrated viral lysates

Enumeration was performed using a BD Influx cell sorter (BD Biosciences) equipped with a 488-nm argon laser and flow meter, as detailed in Zimmerman et al. (4). Cryofrozen samples were thawed, immediately diluted 1:10,000–1:20,000 in 0.02-μm-filtered TE buffer (pH 8.0) and stained in the dark with SYBR Green I nucleic acid dye (0.5X final concentration; Molecular Probes, Inc.) for 15 min at room temperature. The population of viral particles was determined based on green fluorescence (520/35 nm band pass filter, trigger) and Forward Angle Light Scatter as in (4). Samples were run for 2 to 4 min, with the volume determined by weight measurements. Fluorescent polystyrene microsphere standards were added to each sample (0.5 μm Green and 0.75 μm Yellow-Green; Polysciences).

#### Experimental setup

After acclimation of host cultures to the different Fe levels, infection experiments were initiated. On the morning of experiments host cell densities were measured at dawn, 3 hr prior to virus addition, to determine the volume of viral lysate required to achieve a target MOI of 1. Average cell concentrations across both Fe conditions was 3.68 ± 0.13 × 10^6^ cells mL^−1^. The growth rate of the Fe-limited condition on the day of the experiment was 0.40 ± 0.04 day^−1^ and the Fe-replete condition was 0.70 ± 0.03 d^-1^. To ensure an equal number of infectious particles, preliminary MPN assays were conducted, which estimated OlV1 to be 9.1% infective and OlV7 to be 16.8% infective. The viral inoculum represented 1.0 ± 0.1% of the culture volume, and MPN assays were repeated on the inoculum used in the experiment to confirm infectivity and MOI at the time of infection. Triplicate non-infected control cultures for each Fe condition were inoculated with 0.02-μm-filtered TE buffer instead of viral lysate. Following the addition of viruses or buffer, all flasks were manually mixed, and initial samples were collected (t=30 min). Subsequently, culture flasks were sampled every two hours for 24 hours after the addition of viruses. Flow cytometry samples were preserved using glutaraldehyde as described earlier and then stored at -80°C until analysis.

#### Quantification of the concentration of host cells and viral particles over experimental time course

Flow cytometry was used to quantify virions and host cells by thawing glutaraldehyde-preserved samples and running them on the InFlux to enumerate *Ostreococcus* based on FALS (trigger) and red chlorophyll autofluorescence (692/40 band pass filter). Concurrently, the samples were diluted 1:1000 to 1:10,000 in 0.02-μm-filtered TE buffer, stained with SYBR Green I, and analyzed as for viral particles above. Flow cytometry data were analyzed using WinList (Verity Software House).

#### Calculations

Because one step experiments cannot be performed with this eukaryotic phytoplankton species, approaches to understanding temporal dynamics of virus and host populations varied from prokaryote/phage experiments (below). The log2 fold change in abundance was determined at each time point, which corresponds to the number of generations (n) during exponential growth. The log2 fold change was computed using the following formula: *n* _*t*_ = (ln(*N*_*t*_) − ln(*N*_*i*_))/*ln*(2) where *N*_*t*_ is the number of viral particles or host cells at time *t*, and *N*_*i*_ is the number of viral particles or host cells at the initial time point. The specific host growth rate (μ) was determined as follows: *μ* = ln(*N*_*t*_/*N*_*i*_)/(Δ*t*). The latent period was determined by performing a one-tailed paired T-test between the viral particle numbers at the initial time point and each subsequent time point, in addition to computing for between each adjacent set of time points. The first time point difference with a p-value<0.05 was defined as the latent period (i.e., the point at which changes in viral replication and particle production was evident and significant). The burst size estimation involved comparing viral quantities between the “latent period plus one time point” and the “latent period.” The difference in viral quantities was divided by the difference in host cell quantities, accounting for the presence or absence of viruses. The resulting ratio provided an estimate of the average number of viral particles released per infected cell during a single burst of viral replication. A two-sample unequal variance T-Test was used to determine if there was a significant difference in burst size between the Fe-replete and Fe-limited cultures (*p*-value <0.01).

### Experiments with *Vibrio* and vibriophages

#### Initial culturing and viral stock preparation

The two strains of *Vibrio* used in this experiment, and their paired vibriophages, were previously isolated from the littoral zone water column at Canoe Cove, Nahant, Massachusetts, USA, including the *Vibrio splendidus* strain 10N.222.46.C2 (also known as V36; hsp60 sequence: https://www.ncbi.nlm.nih.gov/nuccore/OK263260.1; genome sequence: https://www.ncbi.nlm.nih.gov/nuccore/MDBT00000000.1) and N4-like podovirus 2.130.O_10N.222.46.C2 (75,797 bp genome; genome sequence: https://www.ncbi.nlm.nih.gov/nuccore/1332562253), and the *Vibrio sp*. strain 10N.261.52.C3 (also known as V40; hsp60 sequence: https://www.ncbi.nlm.nih.gov/nuccore/MF464505.1) and myovirus 1.170.O_10N.261.52.C3 (133,692 bp genome; genome sequence: https://www.ncbi.nlm.nih.gov/nuccore/MG592537.1) (6). Long-term *Vibrio* stocks were stored in 50% glycerol at -80°C. *Vibrio* were grown in a modified version of the Vib-FeL medium used in Westrich et al. (7), differing only in the amount of Fe added to the media. Cell culture and phage propagation (as well as experiments) were performed at ∼23°C.

#### Experimental setup

Cells were grown under high-Fe conditions by adding 5 μM Fe, or under low-Fe conditions by adding 1 μM Fe. To minimize Fe carryover into low-Fe treatments, stocks of both host *Vibrio* and phage were always prepared in media with 1 μM Fe. Fresh phage stocks were prepared by adding 10 μL of the prior stock to 5 mL of host culture, incubating for at least 4 hrs, and filtering through a 0.2 μM acid washed polyethersulfone filter for storage at 4°C in 2 mL tubes (Eppendorf Protein LoBind). *Vibrio* cells were enumerated by serial dilution and plating onto agar containing 58 g L^-1^ Zobell Marine Agar 2216 (HiMedia) and 52 mL L^-1^ glycerol. Phage were enumerated by serial dilution and an assay for plaque forming units (PFU; where each plaque is assumed to indicate the presence of one active viral particle in the original solution, or one single infected cell). Assays for PFU were performed by pipetting 100 μL of axenic host culture, a known volume of phage-containing solution, and 2.5 mL of top agar (39 g L^-1^ Zobell Marine Broth 2216, 52 mL glycerol, and 3 g agar) onto bottom agar. Cell abundances and growth rates were measured by absorption at 595 nm, calibrated to absolute abundances based on comparison to serial dilution. Experiments investigating the effect of Fe on growth and viral production were performed three times, and PFU assays were performed in triplicate for each time point.

#### Quantification of plaque-forming units

Infection dynamics (latent period and burst size) were determined for the Fe-replete and Fe-limited experiments in one-step phage propagation experiments. Host cultures were revived by thawing a frozen stock, which was streaked onto agar, and grown overnight. Cells from a single colony were then inoculated into liquid media with either 1 μM Fe or 5 μM Fe using a sterile polystyrene inoculation loop. Once these cultures were in exponential phase growth, phage was added at a MOI of 0.1 and incubated for 15 min to allow phage adsorption. The experiments were then initiated by diluting cultures several orders of magnitude to a phage concentration of 200 phage mL^-1^, ensuring that no additional phage adsorption could take place. The amount of phage which did not adsorb to cells was determined after dilution by enumerating PFU in an aliquot of the experimental mixture gravity-filtered through a 0.2 μm polyethersulfone (Supor) filter. Subsamples of the experimental media were taken every 20 min and assayed for phage concentration by PFU.

#### Calculations

Latent period was determined as the time until a significant increase in phage abundance across triplicates was observed. Burst size was determined as the average PFU after the burst divided by the PFU at the initial time point, both corrected for the amount of unadsorbed phage present at the beginning of the experiment (in contrast, data presented in Fig. 1 shows total PFU). Growth inhibition in Fe-limited cultures was determined by growing uninfected *Vibrio* hosts before and during the one-step experiments, and monitoring growth by absorbance of the media at 595 nm.

**Figure 1.**
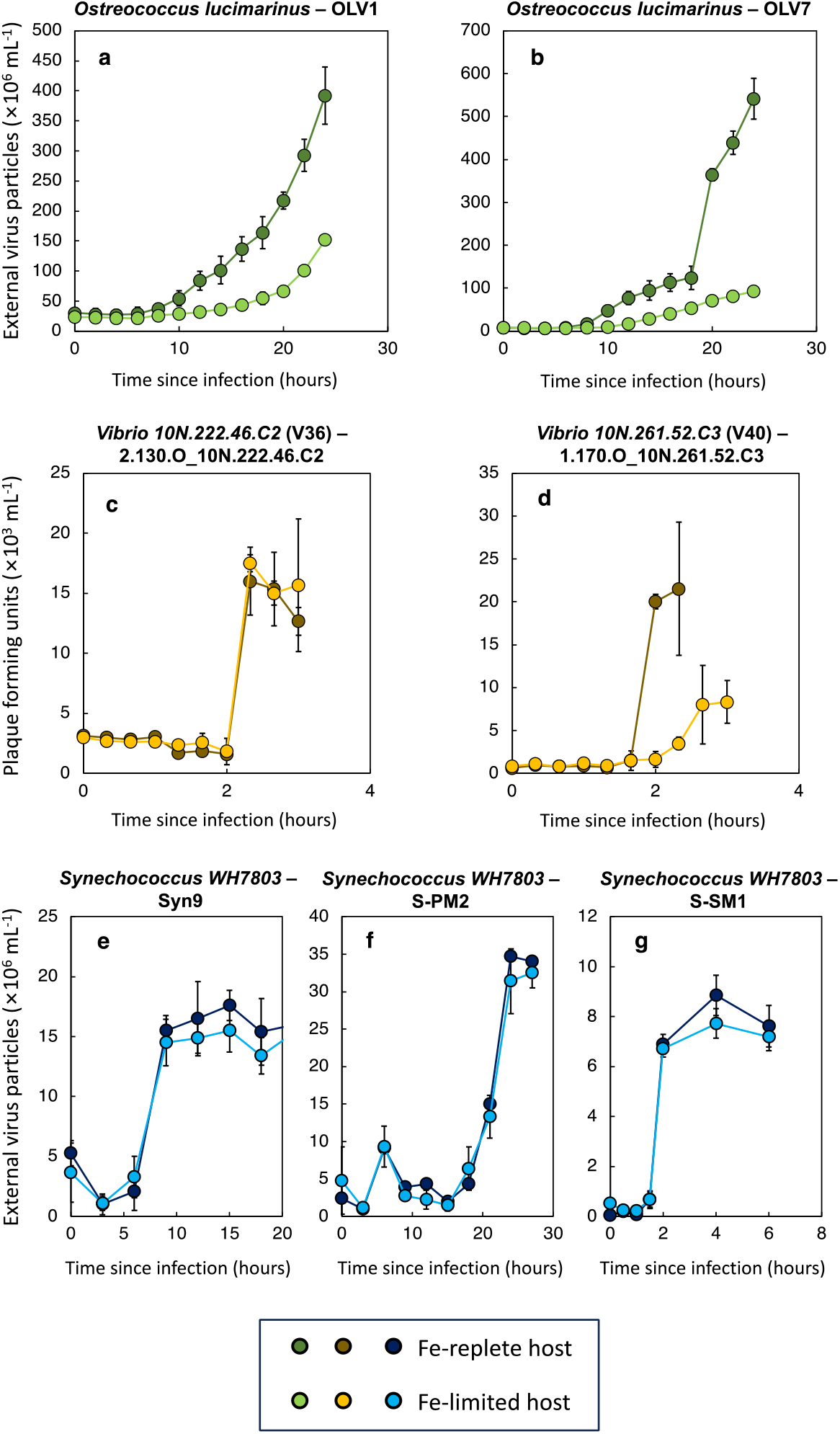
Viral infection dynamics in marine microbes. Infection of the picoeukaryote *Ostreococcus* with the viruses OLV1 (a) and OLV7 (b) resulted in production of fewer viral particles in Fe-limited cultures. One-step viral infection for two *Vibrio* strains (V36 and V40) and their associated viruses demonstrated no impact of Fe on viral infection dynamics of one host-virus pair (c) while in the other case Fe-limitation led to an increase in the latent period of infection and a decrease in burst size (d). One-step viral infection of the cyanobacterium *Synechococcus* with the phages Syn9 (e), S-PM2 (e), and S-SM1 (f) showed no differences in either latent period or burst size between Fe-replete and Fe-limited cultures.

### Experiments with *Synechococcus* and cyanophages

#### Initial culturing and viral stock preparation

Cultures of *Synechococcus* WH7803 (CCMP1334) were grown in a modified SN media (8), using Chelex-purified artificial seawater instead of natural seawater and with either the typical Fe concentrations of 23 μM Fe or reduced concentrations of 230 nM Fe. Cells were grown under a 14:10 light:dark cycle at an irradiance of 40 μmol photons m^−2^ s^−1^ at 23° C, with a bubbled air supply passed through a 0.2 μm polyethersulfone (PES) filter. Phage stocks (myovirus Syn9, 177,300 bp genome, genome sequence: https://www.ncbi.nlm.nih.gov/nuccore/DQ149023.2; myovirus S-PM2, 196,280bp, genome sequence: https://www.ncbi.nlm.nih.gov/nuccore/NC_006820.1; myovirus S-SM1, 174,079bp genome, genome sequence: https://www.ncbi.nlm.nih.gov/nuccore/NC_015282.1) were prepared via propagation on WH7803 host cells grown under low Fe conditions in order to minimize the amount of Fe carried over during inoculation with phage. Viral particles were harvested after cell lysis, by passing the filtrate through a 0.2 μm PES filter to remove cellular debris, and observed by flow cytometry. Phage stocks were then concentrated from 1 L to ∼50 mL using a 100kDa Sartorius 50R to achieve final concentrations of 10^8^ to 10^9^ viral particles mL^-1^ and stored at 4°C until experiment initiation. Concentrations of active phage in stocks and during experiments were determined by most probable number (MPN) assays in 96 well plates by looking for clearing of the *Synechococcus* host. During all experiments, *Synechococcus* cell concentrations were determined using a Guava HPL flow cytometer.

#### Experimental setup

Experiments were initiated by harvesting exponentially growing *Synechococcus* at a cell density of 10^7^ cells mL^-1^. These cells were then concentrated by centrifugation in 50 mL tubes at 5000 x g for 10 min. Cells were then resuspended in media to a concentration of 10^8^ cells mL^-1^. Host and phage were mixed together, using 500 μL of concentrated cells and 500 μL of phage stock at a concentration of 10^7^ viral particles mL^-1^, yielding a multiplicity of infection (MOI) of 0.1. This mixture was incubated for 15 min to allow phage to adsorb to the host. Then 500 μL of the phage-cell mixture was diluted 100-fold into fresh media to initiate the experiment. To ensure that there was no Fe contamination during the concentration process of phage and host, replicate experiments were conducted using heat-killed phage (killed by placing tubes in an 80°C water bath for 15 min). Growth of the host in High Fe and Low Fe media, with the addition of heat-killed phage, was monitored to ensure that the host’s growth was still limited by low Fe concentrations.

#### Quantification of free viral particles

Quantification was performed throughout the experiment using quantitative real-time polymerase chain reaction (qPCR). Primers for Syn9, S-SM1, and S-PM2 were designed using Primer3 software with an ideal primer size of 18-22bp and a product size of ∼120bp. The primer sequences for each phage are as follows: Syn9 forward AGCGATTAAAGCAGTCAACC, Syn9 reverse AGGGAGATTACCAACGTCAA, S-SM1 forward GTCCAGAAGAACTGCGTGGT, S-SM1 reverse GCAATTTTCATGCCCTGATT (Zeng and Chisholm 2012), S-PM2 forward CTACACTTCCAGGCGGTCAG, and S-PM2 reverse TCGAAGGATCTCCGTGGACT (this study). The qPCR response was first calibrated using phage standards with concentrations ranging from 10^4^ to 10^8^ phage mL-1, where the phage concentration was determined by wet-mount enumeration of stock lysate (Cunningham et al. 2015). Prior to analysis, both standards and lysates from the one-step growth curve were diluted 50-fold in 10 mM Tris. A mixture of 10 μL of this diluted sample, 12.5 μL of master mix (iTaq Universal SYBER Green Supermix), and 1.25 μL of both primers was prepared in a 96-well PCR plate. The thermal cycling and measurement of SYBR Green fluorescence was accomplished on a Bio-Rad CFX96 Real-Time PCR Detection System with a 10 min pre-incubation at 95°C, amplification for 39 cycles consisting of denaturation at 95°C for 15 s, annealing at 56°C for 15 s, and extension at 72°C for 30 s, and a melting curve analysis including heating to 95°C for 10 s, annealing at 65°C for 5 s, and extension at 95°C for 5 s.

#### Calculations

The latent period was determined as the time until increase in phage concentration. Burst-size was calculated based on the external phage concentration measured by qPCR after burst, divided by the initial amount of active phage added.

## Notes

### Competing Interest Statement

The authors have declared no competing interest.

